# Super Enhancer-driven LncRNA UNC5B-AS1 Inhibits Inflammatory Phenotypic Transition in Smooth Muscle Cells via Lactylation Modification

**DOI:** 10.1101/2024.05.07.593065

**Authors:** Xiangrui Zhu, Xiangming Pang, Xiaoying Wang, Xiaoyu Guan, Zhaosi Wang, Lixin Zhang, Xiaodong Zheng, Fei Li, Jian Mei, Langlin Ou, Yuxiang Liu, Zitong Meng, Yingli Chen, Cui Ma

## Abstract

**Background:** The phenotypic transition of pulmonary artery smooth muscle cells (PASMCs) is a central pathological alteration in pulmonary artery remodeling that contributes to pulmonary hypertension (PH). The specific role of long noncoding RNAs (lncRNAs), especially super enhancer (SE)-driven lncRNAs, in the phenotypic transformation of PASMCs induced by hypoxia remains unclear. The objective of this study is to determine the mechanistic role of the super enhancer-driven LncRNA UNC5B-AS1 in phenotypic transformation of PASMCs and hypoxia-induced vascular remodeling.

**Methods:** The lncRNA UNC5B-AS1 regulated by super-enhancers was identified in hypoxic human PASMCs through RNA sequencing and H3K27ac ChIP sequencing. Overexpression of lncRNA UNC5B-AS1 in human PASMCs was performed to elucidate its role in the pathogenesis of PH. A conserved functional fragment of lncRNA UNC5B-AS1 was used for the treatment of mouse PH.

**Results:** In PASMCs, we have identified a SE-driven lncRNA called UNC5B-AS1 that regulates phenotype transition. It is transcriptionally activated by the transcription factor FOXP3. We demonstrate that UNC5B-AS1, as a molecular scaffold in mitochondria, stabilizes the interaction between LRPPRC, which is rich in leucine, and oxidative respiratory chain complex IV. This complex regulates the lactylation modification level of upstream promoter regions of inflammatory genes IL-1β, IL-6, and TNF-α by influencing the glycolytic pathway in PASMCs under hypoxic conditions, ultimately affecting the inflammatory phenotype transition of PASMCs.

**Conclusions:** Our findings identify the SE-driven lncRNA UNC5B-AS1 as a novel regulatory factor in hypoxia-induced phenotypic switch of PASMCs, and suggest that overexpression of UNC5B-AS1 may represent a promising therapeutic strategy for PH.

**Highlights:** The long non-coding RNA UNC5B-AS1, regulated by super-enhancers (SE), is downregulated in hypoxic PASMCs, and regulates PASMC phenotype transition through glycolysis.

The super-enhancer region of lncRNA UNC5B-AS1 recruits the transcription factor FOXP3 to its promoter region, forming a chromatin loop to regulate its expression.

The lncRNA UNC5B-AS1 acts as a molecular scaffold stabilizing the interaction between LRPPRC and mitochondrial complex IV.

The lactate generated by glycolysis activation induces lactylation modification of histones at the IL-1β, IL-6, and TNF-α promoters, leading to the inflammatory phenotype transition of PASMCs.

Overexpression of lncRNA UNC5B-AS1 conserved fragment reversed SuHx-induced PH in mice.

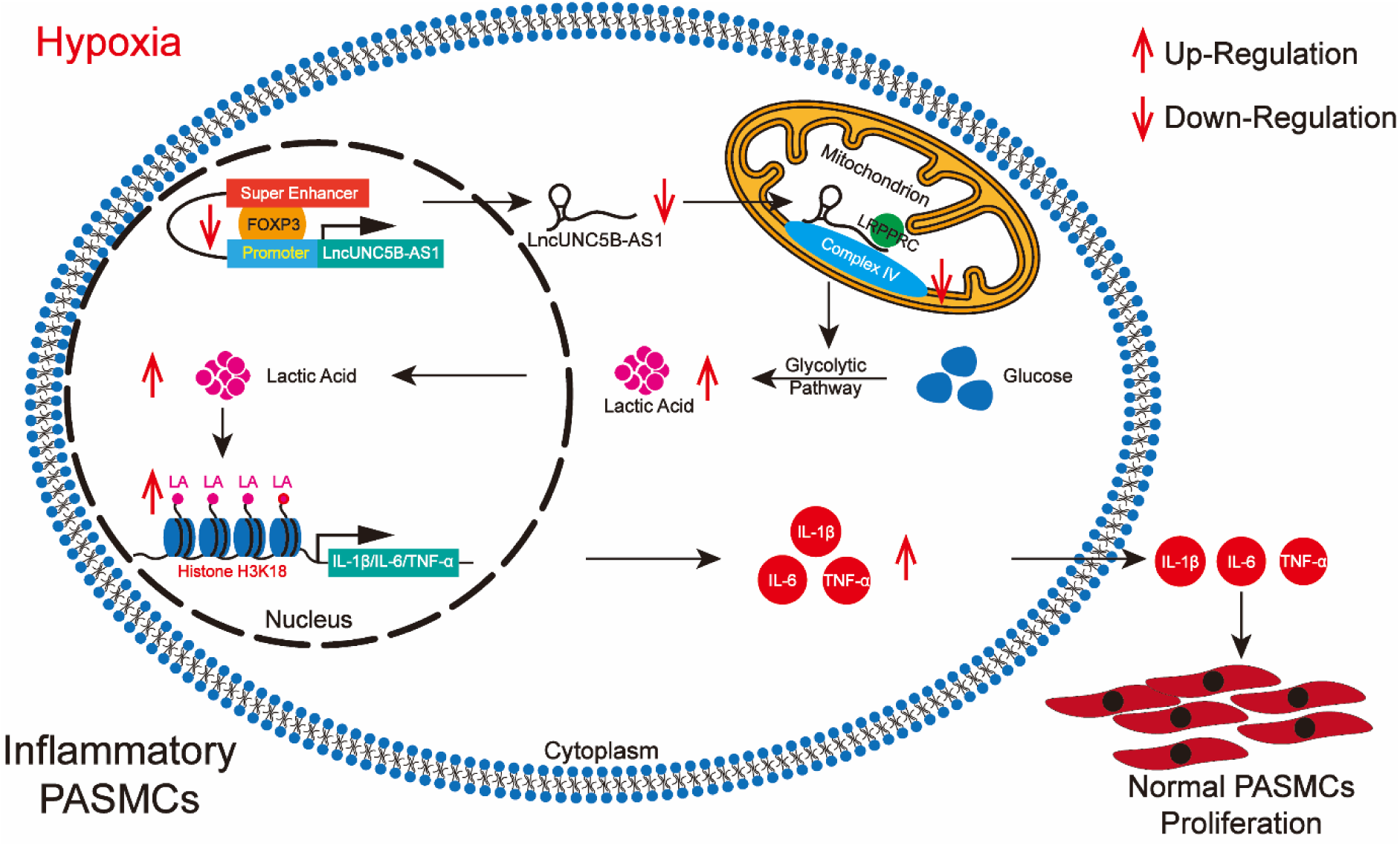

## Introduction

The pathology of pulmonary hypertension (PH) is complex and associated with cardiac, pulmonary, and systemic diseases that contribute to morbidity and mortality^1^. PH is characterized by inflammation, pulmonary vascular remodeling, and pulmonary vascular embolism leading to elevated pulmonary arterial pressure and resistance, right ventricular dysfunction, left ventricular compression, and subsequent heart failure^2,3^. The remodeling process occurs throughout the pulmonary arterial vasculature and involves medial thickening, which is driven by increased proliferation and hypertrophy of pulmonary artery smooth muscle cells (PASMCs), its principal cellular component^4^. However, there is heterogeneity among PASMCs, which can exhibit markedly different changes in proliferative, inflammatory, and extracellular matrix production during remodeling^5–7^. Typically, PASMCs express high levels of contractile proteins such as α-smooth muscle actin (α-SMA), calponin, SM22α and other differentiation markers, which is accompanied by a low rate of proliferation^5,8,9^. Under pathological conditions, PASMCs acquire inflammatory cell markers and release inflammatory cytokines, such as interleukin (IL)-1β, IL-6 and tumor necrosis factor (TNF)-α, contributing to the proinflammatory environment, which is followed by vessel injury and remodeling^10^. These findings suggest that the inflammatory phenotypic transition of PASMCs can contribute to the development of PH.

Long noncoding RNAs (lncRNAs) are non-protein-coding transcripts with lengths greater than 200 nucleotides^11^. An increasing number of studies have shown that lncRNAs play important roles in diverse cellular processes, including proliferation^12^, apoptosis^13^, migration^14^, invasion^15^, chromatin remodeling^16^, metabolism^17^, and immune escape^18^. Super enhancers (SEs) are a widespread class of chromatin regulatory elements in the genome that are closely associated with the expression levels of specific genes^19^. LncRNAs driven by SEs are lncRNAs expressed near SEs that regulate gene expression through interactions with SEs. In gliomas, the SE-driven lncRNA TMEM44-AS1 promotes tumor cell growth through the formation of a positive feedback loop with MYC^20^. The lncRNA NEAT1 has been identified as an SE-driven lncRNA that controls the lineage fate of bone marrow mesenchymal stem cells (BMSCs) during skeletal aging by affecting mitochondrial function^21^. Moreover, lncRNA CARMN has been confirmed to be driven by an SE in vascular smooth muscle cells and is involved in maintaining the contractile phenotype of smooth muscle cells^22^. However, the mechanism of SE-driven lncRNAs in PASMCs inflammatory phenotypic transition and PH development needs to be further examined.

Emerging evidence shows that lactate has regulatory functions in physiologic and pathologic cells and induces dramatic changes in gene expression, suggesting that lactate is not simply a waste product of glycolysis^23^. In response to hypoxia, cells undergo metabolic reprogramming by inhibiting oxidative phosphorylation and enhancing glycolysis, thereby stimulating the production of lactate^24^. Recent studies indicated that hypoxia induced intracellular lactate production and increased histone Kla but not Kac levels in MCF-7 cells^23^. Lactate-derived lactylation of histone lysine residues is an epigenetic modification that directly stimulates gene transcription^25^. Histone lactylation modification can influence the transition of inflammatory macrophage phenotypes, thereby impacting inflammation levels and participating in the progression of related diseases^25,26^. Thus, controlling the glycolytic switch via lactylation suggests novel therapeutic opportunities to treat PH.

In this study, we showed for the first time that lncRNA UNC5B antisense RNA 1(UNC5B-AS1, Chr10q22.1) was regulated by SEs in PASMCs. It is transcriptionally activated by the transcription factor forkhead box protein P3 (FOXP3). Our data indicated that UNC5B-AS1 served as a molecular scaffold in mitochondria and interacted with leucine-rich PPR motif-containing protein (LRPPRC) and mitochondrial complex IV. Under hypoxic conditions, the loss of UNC5B-AS1 expression impaired mitochondrial oxidative phosphorylation, leading to abnormal activation of glycolysis. The accumulation of lactate due to the activation of glycolysis in hypoxia, affected the lactylation of inflammatory factor promoters in PASMCs and ultimately promoted the transition of PASMCs to an inflammatory phenotype and contributed to the progression of PH.

## Methods

### Animal

This study used adult male C57BL/6 mice (6-9 weeks old) obtained from the Experimental Animal Center of the Second Affiliated Hospital of Harbin Medical University. All experimental procedures have obtained approval from the Ethics Committee of Harbin Medical University (HMUDQ20230831001). All animal studies were conducted in compliance with the National Institutes of Health (NIH) guidelines for the care and use of laboratory animals. The RNA cloning constructs, and serotype 5 adenovirus-associated virus (AAV5) were synthesized by GeneChem (China). After randomization, the mice were divided into groups and intranasally infected with 10^11^ genome equivalents of AAV5 vector solution diluted with 20 μL of Hank’s balanced salt after isoflurane anesthesia. After 14 days, the mice were assigned to normoxia (21% O_2_) and hypoxia (10% O_2_) conditions, as described in previous studies^27^. In the Sugen hypoxia model, the mice were injected once per week with the VEGF inhibitor SU5416 (Med Chem Express, USA) at a concentration of 20 mg/kg. Subsequently, the mice were subjected to 3 weeks of hypoxia (10% O_2_) followed by 2 weeks of reoxygenation (21% O_2_). The control group was maintained in a normal oxygen environment, and the oxygen concentration was continuously monitored for 5 weeks using an oxygen analyzer (BioSpherix, USA). After constructing the model, euthanasia was performed on mice using a small animal euthanasia system (Yuyan Instruments, China) by inhalation of 100% CO_2_, followed by cervical dislocation as a secondary confirmation of death. Lung tissue was collected for further experiments. The right ventricular (RV) hypertrophy index (RV/(LV+S)), is calculated by dividing the weight of the right ventricular free wall by the sum of the weight of the septum plus the left ventricular free wall.

### Small Animal Echocardiography and Right Ventricular Systolic Pressure

After the completion of hypoxia and SU5416 treatment, the mice underwent echocardiography using the Vevo2100 imaging system equipped with an 18 to 38MHz transducer probe (VisualSonics, Canada), as described previously^28^. Mice were anesthetized using a desktop anesthesia machine (Harvard Apparatus, USA) with inhalation of a concentration of 1% to 3% isoflurane. The mice were placed on a preheated imaging platform and the respiratory rate was continuously monitored throughout the entire echocardiography process. The pulmonary artery acceleration time (PAAT) and pulmonary artery velocity-time integral (PAVTI) were measured using pulse wave Doppler. Transthoracic echocardiography was performed in the short-axis view at the level of the sternum to measure left ventricular ejection fraction at end-diastole and end-systole. All measurements were performed by experienced researchers, and data analysis was conducted in a blinded manner using Vevo2100 software. Right ventricular systolic pressure was measured using the PowerLab monitoring system (AD Instruments, USA). The pressure catheter (Scisense Inc) was inserted into the right ventricular vein via the superior vena cava, and right ventricular systolic pressure was continuously recorded for 10 to 30 minutes.

### ChIP-seq

ChIP-seq was performed by Guangzhou Epibiotek Co., Ltd. Briefly, the cells were cross-linked with 1% formaldehyde for 10 minutes, followed by quenching the cross-linking reaction with 0.125M glycine. The cells were then frozen on dry ice and stored until further use. After isolating the cell nuclei, the chromatin was sheared to fragment it into sizes ranging from 200 to 500 bp. 5 μg of anti-H3K27ac antibody (Cell Signaling) was incubated overnight with the fragmented chromatin. Protein A magnetic beads were used to capture immune complexes. The chromatin was eluted in reverse cross-linking buffer followed by incubate the tube at 65℃ for 3h. After digestion with RNase and Proteinase K, the fragments were purified and quantified. ChIP DNA was processed for library generation using the QIAseq Ultralow Input Library Kit (QIAGEN) following the manufacturer’s protocol. After library construction, the samples were subjected to sequencing analysis.

### RNA-seq

PASMCs cultured under normoxia/hypoxia conditions were harvest or RNA-seq. Total RNA was isolated using TRIzol reagent (Thermo Fisher Scientific, Massachusetts) according to the manufacturer’s protocol. VAHTS Stranded mRNA-seq Library Prep Kit for Illumina V2 (Vazyme Biotech, NR612-02) was used for library preparation according to the instructions. The libraries were then sequenced on the Illumina NovaSeq 6000 platform by Epibiotek Co., Ltd. in Guangzhou.

### Statistical Analysis

The statistical analysis was performed using GraphPad Prism 8 software. Prior to statistical testing, the data were checked for normality and homogeneity of variances (F test). Student’s t-test (unpaired) was used for analyzing data with equal variances in two groups, while Welch’s corrected t-test was used for analyzing data with unequal variances in two groups. One-way analysis of variance (ANOVA) with Tukey’s post hoc test was used to compare multiple groups with equal variances, while Brown-Forsythe and Welch’s ANOVA with Tamhane T2 post hoc test were used to compare multiple groups with unequal variances. Nonparametric analysis was used for non-normally distributed data, including Mann-Whitney U test or Kruskal-Wallis test for two group analysis, followed by Dunn’s post hoc test for >2 groups. Data are presented as mean ± SEM, and a P value of <0.05 is considered statistically significant.

## Results

### UNC5B-AS1 is a super enhancer-driven lncRNA that is downregulated by hypoxia

To identify differentially expressed lncRNAs in PH, we performed RNA-seq and H3K27ac ChIP-seq assays on PASMCs subjected to normoxia/hypoxia. Our results revealed significant downregulation of UNC5B-AS1 under hypoxic conditions (Fig.1A, B). H3K27ac ChIP-seq analysis was used to identify super enhancers in the upstream regulatory regions of genes, the hockey-stick plot demonstrated increased super enhancer activity upstream of UNC5B-AS1 under normoxic conditions (Fig.1C). Similarly, there was significantly increased abundance of H3K27ac modification upstream of the gene in normoxic conditions (Fig.1D). We further validated the significant downregulation of UNC5B-AS1 under hypoxic conditions in PASMCs (Fig.1E), and in response to JQ1 (an effective BET bromodomain inhibitor) treatment, the expression of UNC5B-AS1 was significantly inhibited after the chromatin interaction was suppressed (Fig.1F). Fluorescence in situ hybridization (FISH) indicated cytoplasmic localization of UNC5B-AS1 in PASMCs (Fig.1G). Taken together, these findings suggest that UNC5B-AS1 is regulated by super enhancers in PASMCs and that its expression is downregulated and localized in the cytoplasm under hypoxic conditions.

**Fig. 1.**
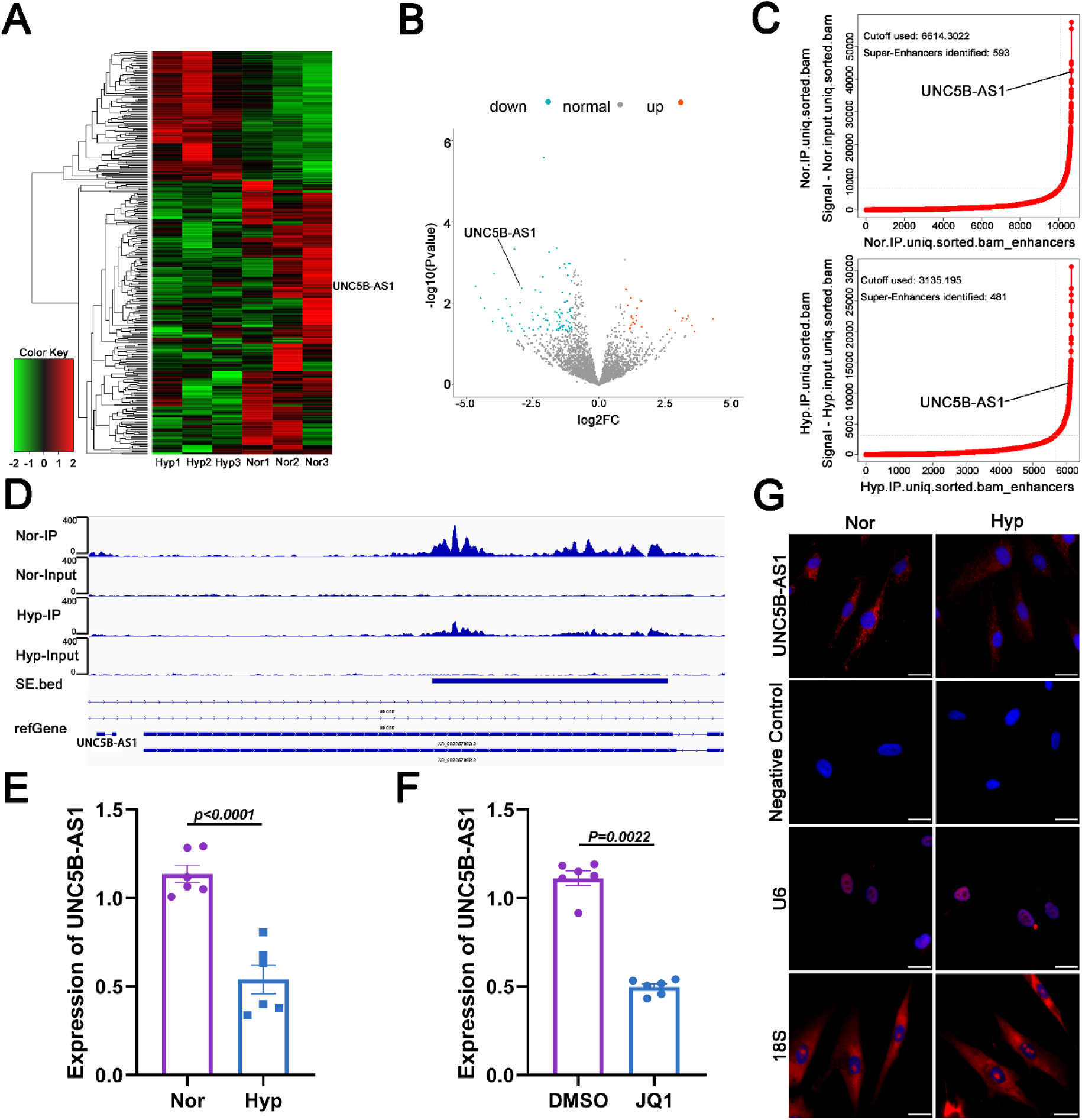
The super enhancer-driven lncRNA UNC5B-AS1 is downregulated in hypoxic PASMCs. **A** The heatmap displays the differentially expressed genes identified in normoxic/hypoxic PASMCs based on RNA-seq analysis (n=3). **B** The RNA-seq results are visualized as a volcano plot, displaying the expression pattern of UNC-5B-AS1 in normoxic/hypoxic PASMCs. **C** The hockey stick plot displays the activity of super enhancers and enhancers upstream of UNC5B-AS1 in normoxic/hypoxic PASMCs (Super enhancers are defined as slope >1, while enhancers slope <1). **D** To visualize the enrichment profile of H3K27ac signal peaks upstream of UNC5B-AS1 in normoxic/hypoxic PASMCs by using IGV (Integrative Genomics Viewer). **E** The expression levels of UNC5B-AS1 in normoxic/hypoxic PASMCs (n=6). **F** The expression levels of UNC5B-AS1 after treatment of PASMCs with JQ1 (n=6). Statistical comparison was performed by the paired Student’s t test (two-sided). **G** FISH results for the localization of UNC5B-AS1 in normoxic/hypoxic PASMCs. Scale bars=50μm. All data are presented as mean ± SEM. Nor, normoxia and Hyp, hypoxia.

### FOXP3 regulates the expression of UNC5B-AS1 through chromatin remodeling

UNC5B-AS1, which is a super enhancer-regulated lncRNA that is downregulated under hypoxic conditions, motivated us to further investigate the upstream regulatory mechanisms driving its expression. We used three bioinformatics tools to screen for potential transcription factors that could regulate UNC5B-AS1 expression (Fig.S1A left). Based on the number and binding agent of the binding sites within the upstream promoter region of UNC5B-AS1, we identified FOXP3 as a key transcription factor that regulates UNC5B-AS1 expression (Fig.S1A right). Immunoblotting showed that FOXP3 was downregulated under hypoxic conditions (Fig.2A). Subsequently, based on the H3K27ac ChIP-seq results, we divided the upstream super enhancer region of UNC5B-AS1 into three segments (E1-E3, Fig.2B). ChIP-qPCR experiments using H3K27ac (Histone 3 Lysine 27 Acetylation, super-enhancer marker), H3K4me1 (Histone 3 Lysine 4 Monomethylation, enhancer marker), and FOXP3 antibodies revealed that FOXP3 had a strong binding affinity to segment E2 of the UNC5B-AS1 upstream super enhancer region (Fig.2C). Moreover, we divided the average span of the UNC5B-AS1 upstream promoter region into three segments (P1-P3, Fig.S1B) and performed ChIP-qPCR experiments by using H3K27ac, H3K4me1, and FOXP3 antibodies. The results demonstrated that FOXP3 has a strong binding affinity to segment P1 of the UNC5B-AS1 upstream promoter region (Fig.2D). Coimmunoprecipitation confirmed the interaction among FOXP3, H3K27ac, and H3K4me1 (Fig.2E). These findings confirmed that FOXP3 was recruited by the upstream super enhancer E2 region and acted on the promoter P1 region of UNC5B-AS1, forming a chromatin loop to regulate the expression of the super enhancer-driven lncRNA UNC5B-AS1.

**Fig. 2.**
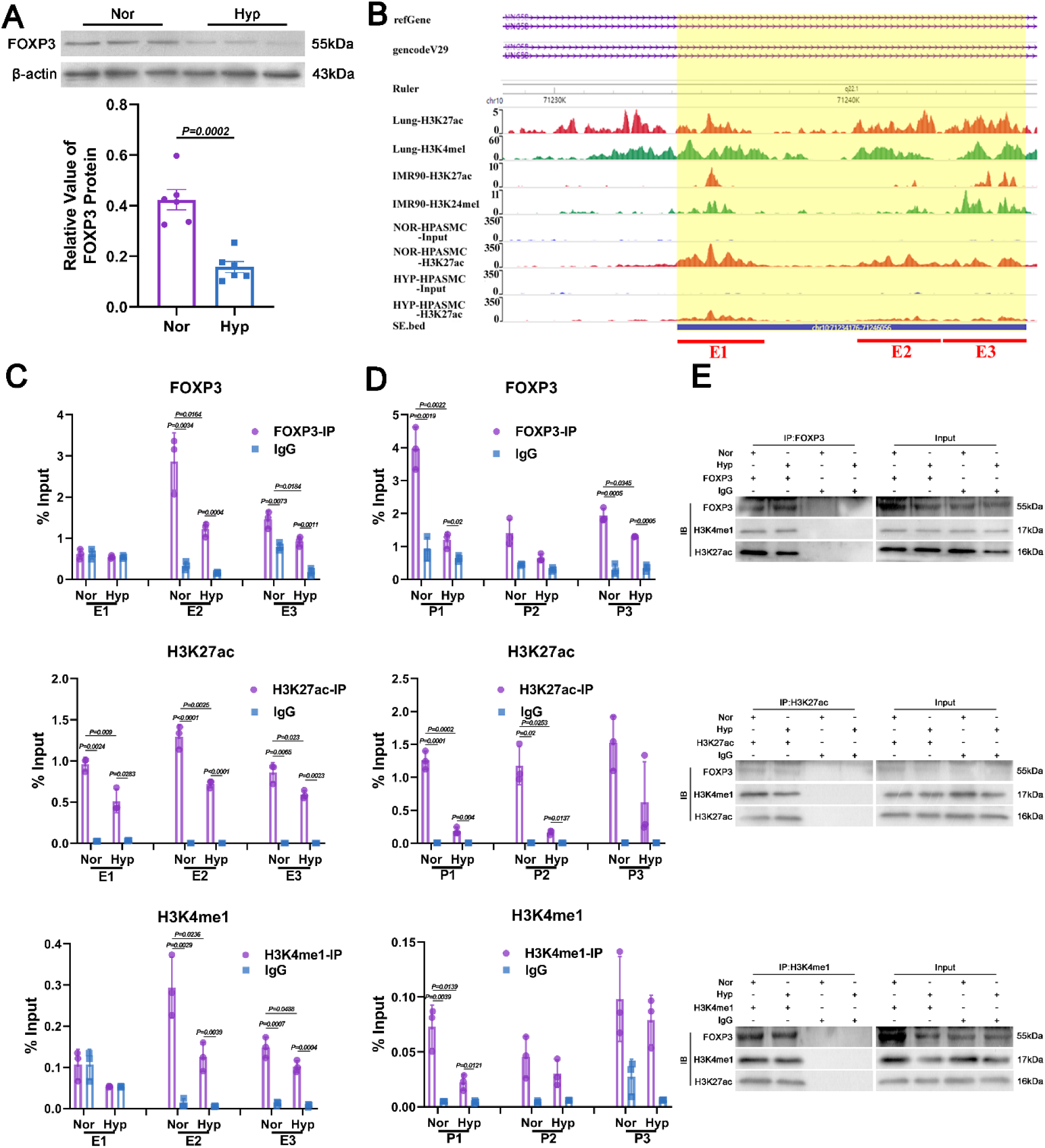
FOXP3 regulates the expression of UNC5B-AS1 through chromatin remodeling. **A** Western blot analysis of FOXP3 in normoxic/hypoxic PASMCs (n=6). **B** Schematic diagram illustrating the segmentation of the upstream super enhancer region of UNC5B-AS1 based on H3K27ac ChIP-seq results. FOXP3, H3K27ac, and H3K4me1 ChIP-qPCR results showing their enrichment in the upstream super enhancer region (**C**) and promoter region (**D**) of UNC5B-AS1 (n=3). **E** Results of co-immunoprecipitation with FOXP3, H3K27ac, and H3K4me1. Statistical comparison was performed by the paired Student’s t test (two-sided). All data are presented as mean ± SEM. Nor, normoxia, Hyp, hypoxia, P, promoter, E, super enhancer, IP, immunoprecipitation, IB, Immunoblotting and IgG, Immunoglobulin G.

### UNC5B-AS1 inhibits hypoxia-induced inflammatory phenotypic transition in PASMCs

To investigate the role of UNC5B-AS1 in hypoxic PH, we constructed an overexpression plasmid for UNC5B-AS1 and validated its overexpression efficiency by RT-qPCR (Fig.S2A). Subsequently, we overexpressed UNC5B-AS1 under hypoxic conditions and found that its overexpression significantly inhibited the transition of hypoxia-induced PASMCs from a contractile phenotype to a synthetic phenotype (Fig.3A, B and Fig.S2B). Inflammatory smooth muscle cells actively secrete inflammatory cytokines such as IL-1β, IL-6, and TNF-α^29^. UNC5B-AS1 overexpression notably attenuated the hypoxia-induced inflammatory phenotypic transition in PASMCs by suppressing the expression of IL-1β, IL-6, and TNF-α (Fig.3C and Fig.S2C). These results confirm that UNC5B-AS1 alleviates the hypoxia-induced inflammatory phenotypic transition in PASMCs.

**Fig. 3.**
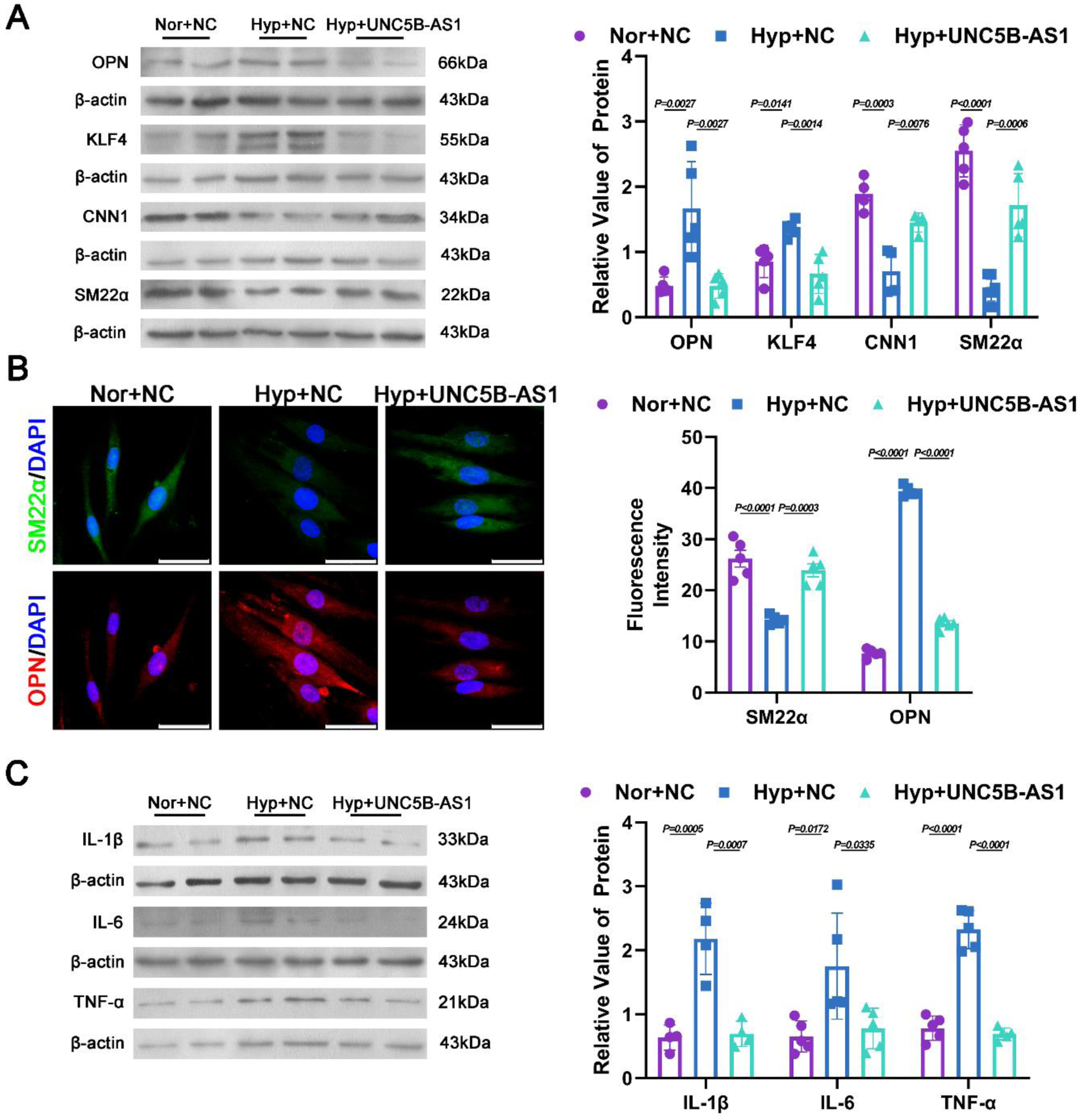
UNC5B-AS1 alleviates hypoxia-induced inflammatory phenotypic transition in PASMCs. **A** Western blot analysis of OPN, KLF4, CNN1 and SM22α in PASMCs (n=5 for OPN, KLF4 and SM22α, n=4 for CNN1). **B** Representative immunofluorescence images of PASMCs stained with anti-SM22α (green) antibody, anti-OPN (red) antibody and DAPI (blue). Scale bars=100 μm. **C** Western blot analysis of IL-1β, IL-6 and TNF-α in PASMCs (n=5 for IL-6 and TNF-α, n=4 for IL-1β). All data are presented as mean ± SEM. One-way ANOVA with Tukey post hoc test was used to compare multiple groups with equal variance, and Brown-Forsythe and Welch ANOVA with Tamhane T2 post hoc test was used to compare multiple groups with unequal variance. Nor, normoxia, Hyp, hypoxia, NC, Negative control and UNC5B-AS1, UNC5B-AS1 overexpression.

### UNC5B-AS1 inhibits PASMCs proliferation by modulating inflammatory responses

Smooth muscle cells (SMCs) undergo phenotypic transition toward an inflammatory phenotype and release inflammatory cytokines to participate in the perivascular inflammatory microenvironment, ultimately leading to vascular remodeling^30^. To investigate the crucial role of inflammatory PASMCs in the development of PH, we constructed a coculture system to simulate the inflammatory microenvironment around blood vessels. First, PASMCs were cultured under different experimental conditions for 24 hours. The cell culture supernatant (conditioned medium, CM) was collected by centrifugation, and the levels of IL-1β, IL-6, and TNF-α were measured. A portion of the remaining conditioned medium was mixed with normal cell culture medium at a 1:1 ratio and used to stimulate normal PASMCs. Proliferation-related markers were assessed after 24 hours to evaluate the impact of the inflammatory microenvironment on normal PASMCs proliferation (Fig.S3A). Our results demonstrated that UNC5B-AS1 overexpression significantly suppressed the secretion of IL-1β, IL-6, and TNF-α in hypoxia-induced PASMCs (Fig.S3B). Conditioned media from UNC5B-AS1-overexpressing cells significantly inhibited the proliferation (Fig.S3C) of normal PASMCs under hypoxic conditions, which was accompanied by reduced cell viability (Fig.S3D) and decreased expression of PCNA (Fig.S3E). These findings suggest that UNC5B-AS1 can regulate the inflammatory microenvironment around blood vessels by suppressing the inflammatory phenotypic transition of PASMCs, thereby alleviating hypoxia-induced PASMCs proliferation.

### UNC5B-AS1 regulates the inflammatory microenvironment in hypoxia through lactylation

The subcellular localization of lncRNAs determines their functions. We further examined the subcellular localization of UNC5B-AS1. By using TOMM20 as a marker of mitochondria and performing costaining with FISH, we found strong colocalization of UNC5B-AS1 with mitochondria (Fig.4A). Subsequently, we isolated cytoplasm and mitochondria, RT-qPCR showed the presence of UNC5B-AS1 in both compartments, and there was decreased expression under hypoxic conditions (Fig.4B). Mitochondria, which are central organelles involved in cellular energy metabolism, are closely related to glucose metabolism. Based on the subcellular localization of UNC5B-AS1 and abnormal glucose metabolism under hypoxic conditions, we further assessed the impact of UNC5B-AS1 on glycolysis, as in our previous study^31^. Immunoblotting showed that UNC5B-AS1 overexpression inhibited the upregulation of HKII and PKM2, which are key enzymes in glycolysis during hypoxia (Fig.4C). Lactate, which is the main product of glycolysis, serves as a substrate for posttranslational modifications of proteins^25^. Treatment of PASMCs with the glycolysis inhibitor 2-DG attenuated the hypoxia-induced increase in protein lactylation and proliferation (Fig.S4A, B). Additionally, using panlactylation antibody, we confirmed that UNC5B-AS1 overexpression abrogated the increase in protein lactylation induced by hypoxia (Fig.4D). Finally, we assessed the H3K18la levels of the promoters of the proinflammatory cytokines IL-1β, IL-6, and TNF-α. ChIP-qPCR confirmed that UNC5B-AS1 modulated the lactylation levels of these gene promoters, thereby affecting their expression (Fig.4E). These data suggest that UNC5B-AS1 affects the lactate modification of the IL-1β, IL-6, and TNF-α promoter regions through the glycolysis pathway, thereby increasing their expression and participating in the inflammatory phenotypic transition of PASMCs.

**Fig. 4.**
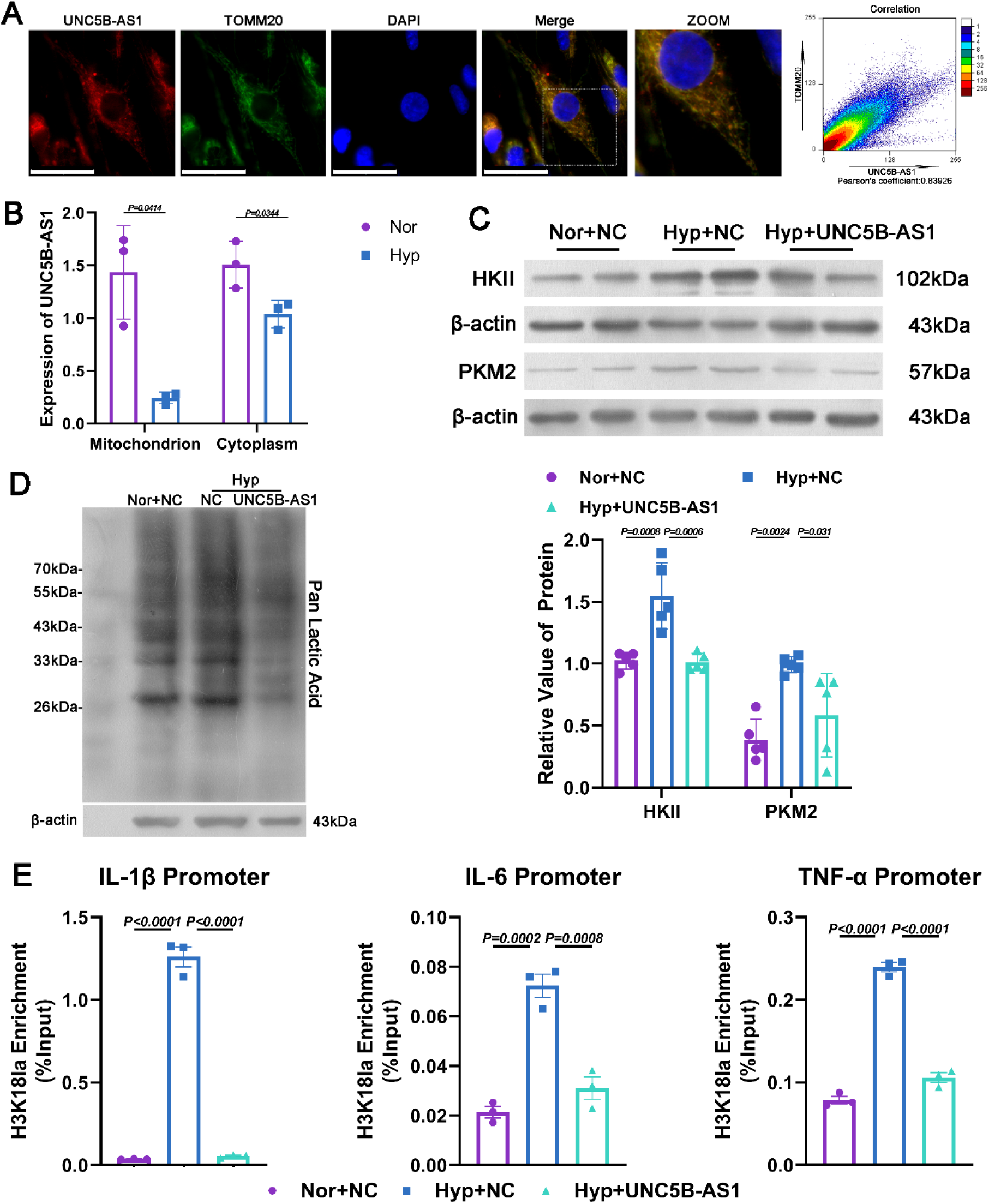
UNC5B-AS1 inhibits the expression of inflammatory factors through lactylation modification. **A** Representative immunofluorescence images of PASMCs stained with anti-TOMM20 (green) antibody, UNC5B-AS1 probe (red) and DAPI (blue), the Pearson correlation coefficient reflects the correlation between two fluorescence labels in co-localization. Scale bars=100 μm. **B** Expression of UNC5B-AS1 in PASMC mitochondrion and cytoplasm (n=3). Statistical comparison was performed by the paired Student’s t test (two-sided). **C** Western blot analysis of the expression of HKII and PKM2 in PASMCs (n=5). **D** Western blot analysis of pan-lactylation modification in PASMCs. **E** ChIP-qPCR analysis of H3K18la modification in IL-1β, IL-6 and TNF-α promoter (n=3). All data are presented as mean ± SEM. One-way ANOVA with Tukey post hoc test was used to compare multiple groups with equal variance, and Brown-Forsythe and Welch ANOVA with Tamhane T2 post hoc test was used to compare multiple groups with unequal variance. Nor, normoxia, Hyp, hypoxia, NC, Negative control and UNC5B-AS1, UNC5B-AS1 overexpression.

### UNC5B-AS1 acts as a molecular scaffold to modulate the activity of mitochondrial complex IV through LRPPRC

To further explore the specific mechanism by which UNC5B-AS1 influences the glycolytic pathway, we used three bioinformatics websites (catRAPID, RNAct, and RNAInter) and the subcellular localization of UNC5B-AS1 to identify LRPPRC as a downstream mitochondrial-related protein associated with UNC5B-AS1 (Fig.S5A). Immunofluorescence staining demonstrated that LRPPRC was localized within mitochondria, confirming its role as a mitochondrial-related protein (Fig.S5B). The colocalization of UNC5B-AS1 and LRPPRC was confirmed by FISH and immunofluorescence staining (Fig.5A). Immunoblotting indicated that LRPPRC was downregulated under hypoxic conditions and that UNC5B-AS1 did not influence its expression (Fig.5B). RNA Pull-down assays (Fig.5C) and RIP experiments (Fig.5D) demonstrated the interaction between UNC5B-AS1 and LRPPRC. Based on these findings, we confirmed the interaction between UNC5B-AS1 and LRPPRC, both of which are associated with mitochondria, and the expression of LRPPRC was not affected by UNC5B-AS1.

**Fig. 5.**
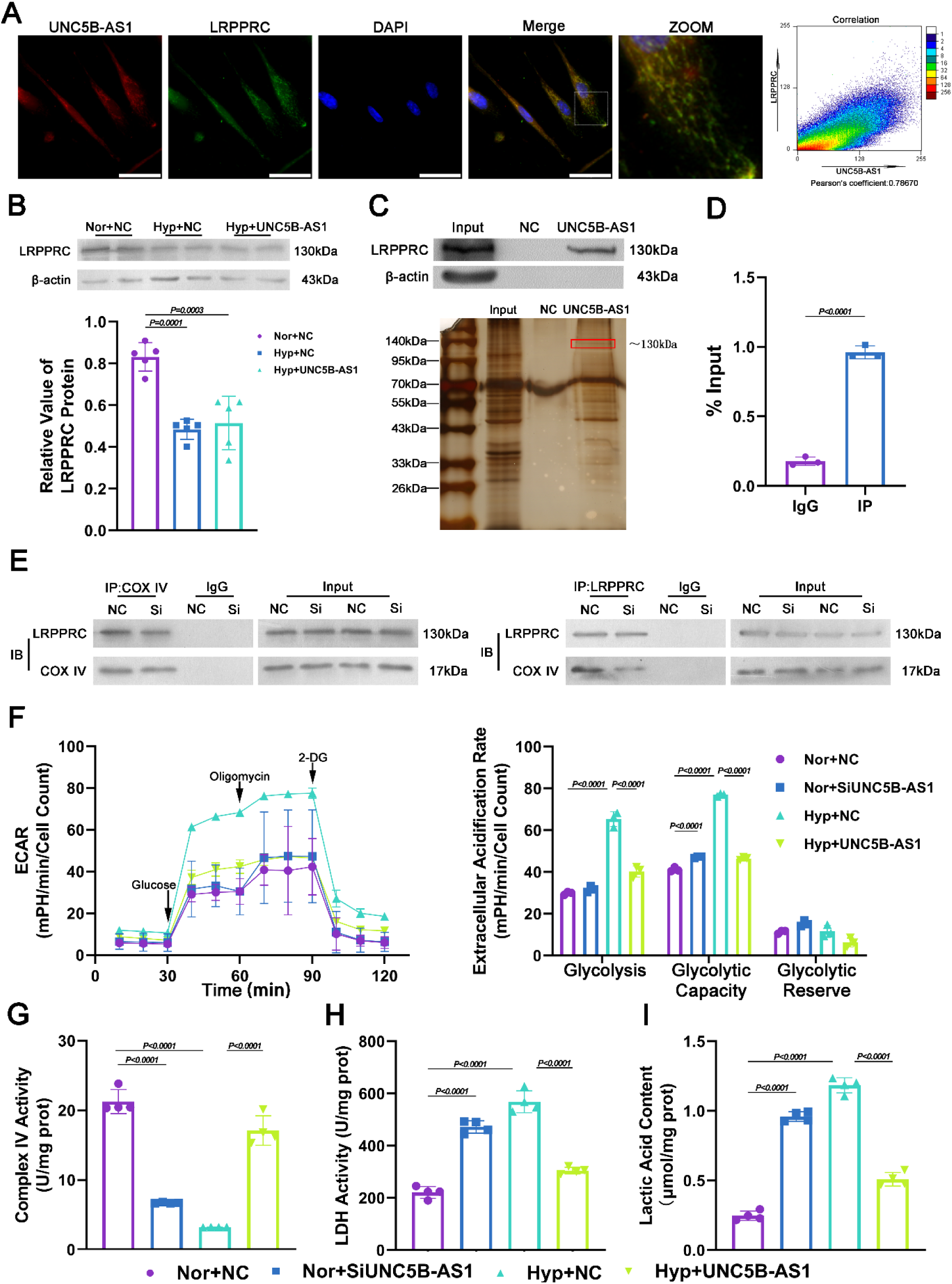
UNC5B-AS1 acts as a molecular scaffold by binding to LRPPRC, influencing the activity of mitochondrial complex IV. **A** Representative immunofluorescence images of PASMCs stained with anti-LRPPRC (green) antibody, UNC5B-AS1 probe (red) and DAPI (blue), the Pearson correlation coefficient reflects the correlation between two fluorescence labels in co-localization. Scale bars=100 μm. **B** Western blot analysis of LRPPRC in PASMCs (n=5). **C** Up panel, RNA pull-down analysis was performed to examine the binding between UNC5B-AS1 and LRPPRC in PASMCs. Below panel, silver staining results of RNA Pull-down in PASMCs. **D** RIP analysis was performed to examine the binding between UNC5B-AS1 and LRPPRC in PASMCs (n=3). Statistical comparison was performed by the paired Student’s t test (two-sided). **E** Results of co-immunoprecipitation with mitochondrial complex IV (COX IV) and LRPPRC after siUNC5B-AS1. **F** Glycolysis and glycolytic reserve following treatment with 10 mmol/L glucose and 1 µM oligomycin were measured in cultured PASMCs, and nonglycolytic acidification after treatment with 100 mmol/L 2-deoxyglucose was detected (n=3). Mitochondrial complex IV activity (**G**), LDH activity (**H**) and intracellular lactic acid contents (**I**) in PASMCs (n=4). All data are presented as mean ± SEM. One-way ANOVA with Tukey post hoc test was used to compare multiple groups with equal variance, and Brown-Forsythe and Welch ANOVA with Tamhane T2 post hoc test was used to compare multiple groups with unequal variance. Nor, normoxia, Hyp, hypoxia, NC, Negative control, SiUNC5B-AS1, UNC5B-AS1 small interfering RNA and UNC5B-AS1, UNC5B-AS1 overexpression.

LRPPRC was reported to exert its biological effects by influencing the activity of mitochondrial complex IV^32^. We propose that UNC5B-AS1 is a molecular scaffold between LRPPRC and mitochondrial complex IV, facilitating their interaction and maintaining the activity of mitochondrial complex IV. First, we designed small interfering RNAs (siRNA) targeting UNC5B-AS1 and confirmed the efficiency of UNC5B-AS1 knockdown in PASMCs (Fig.S5C). Immunoprecipitation showed that knockdown of UNC5B-AS1 affected the interaction between mitochondrial complex IV and LRPPRC (Fig.5E). Subsequently, we performed real-time monitoring of glycolysis using the Seahorse XFe24 extracellular flux analyzer. As expected, knockout of UNC5B-AS1 activated glycolysis in PASMCs under normoxic conditions, while overexpression of UNC5B-AS1 decreased glycolysis under hypoxia-induced stress (Fig.5F). Next, we evaluated the impact of UNC5B-AS1 on the activity of mitochondrial complex IV, and the results showed that knockdown of UNC5B-AS1 under normoxic conditions impaired the activity of mitochondrial complex IV, while UNC5B-AS1 overexpression alleviated the hypoxia-induced decrease in the activity of mitochondrial complex IV (Fig.5G). Finally, we assessed the effects of UNC5B-AS1 on lactate dehydrogenase (LDH, catalyzing the conversion of pyruvate to lactate) activity and intracellular lactate levels. The results confirmed that UNC5B-AS1 overexpression inhibited the hypoxia-induced increase in LDH activity and intracellular lactate levels, while knockdown of UNC5B-AS1 under normoxic conditions increased LDH activity (Fig.5H) and intracellular lactate levels (Fig.5I). These findings confirm that UNC5B-AS1 acts as a molecular scaffold between LRPPRC and mitochondrial complex IV, thereby regulating mitochondrial metabolism and intracellular lactate production.

### The UNC5B-AS1 conserved fragment alleviates hypoxia-induced pulmonary hypertension in vivo

To determine whether UNC5B-AS1 is functionally important in PH in vivo, we first used the catRAPID tool to identify the functional fragment of UNC5B-AS1 that bound to LRPPRC (Fig.S6A). We found a conserved sequence of this functional fragment in mice and named it the UNC5B-AS1 conserved fragment (UA1CF, Fig.S6B). Subsequently, we validated the expression of UA1CF in lung tissues of mice with SU5416 combined with hypoxia-induced PH (SuHx mice) by using RT-qPCR and observed the downregulation of UA1CF expression in SuHx mice (Fig.S6C). RIP experiments on mouse PASMCs confirmed the binding of UA1CF with LRPPRC (Fig.S6D). Next, we constructed an adenovirus overexpressing UA1CF and infected SuHx mice (Fig.6A). After hypoxic treatment, we assessed the impact of adenoviral infection on mice by evaluating body weight and found that there was no significant difference in body weight among the groups (Fig.S6E). RT-qPCR confirmed the successful expression of UA1CF in mouse lung tissues (Fig.6B). We then evaluated SuHx-related PH indicators in mice and found that UA1CF overexpression alleviated the increase in right ventricular hypertrophy (Fig.6C) and right ventricular systolic pressure (RVSP, Fig.6D, E), as well as the reduction in pulmonary artery acceleration time (PAAT, Fig.6F) and pulmonary artery velocity-time integral (PAVTI, Fig.6G) in SuHx mice. However, it did not affect left heart function in mice (Fig.6H). Histological analysis (H&E staining and Masson’s trichrome staining) revealed that UA1CF overexpression alleviated pulmonary arterial thickening and perivascular fibrosis in SuHx mice (Fig.6I). These results confirm that UA1CF can alleviate the symptoms of PH in SuHx mice.

**Fig. 6.**
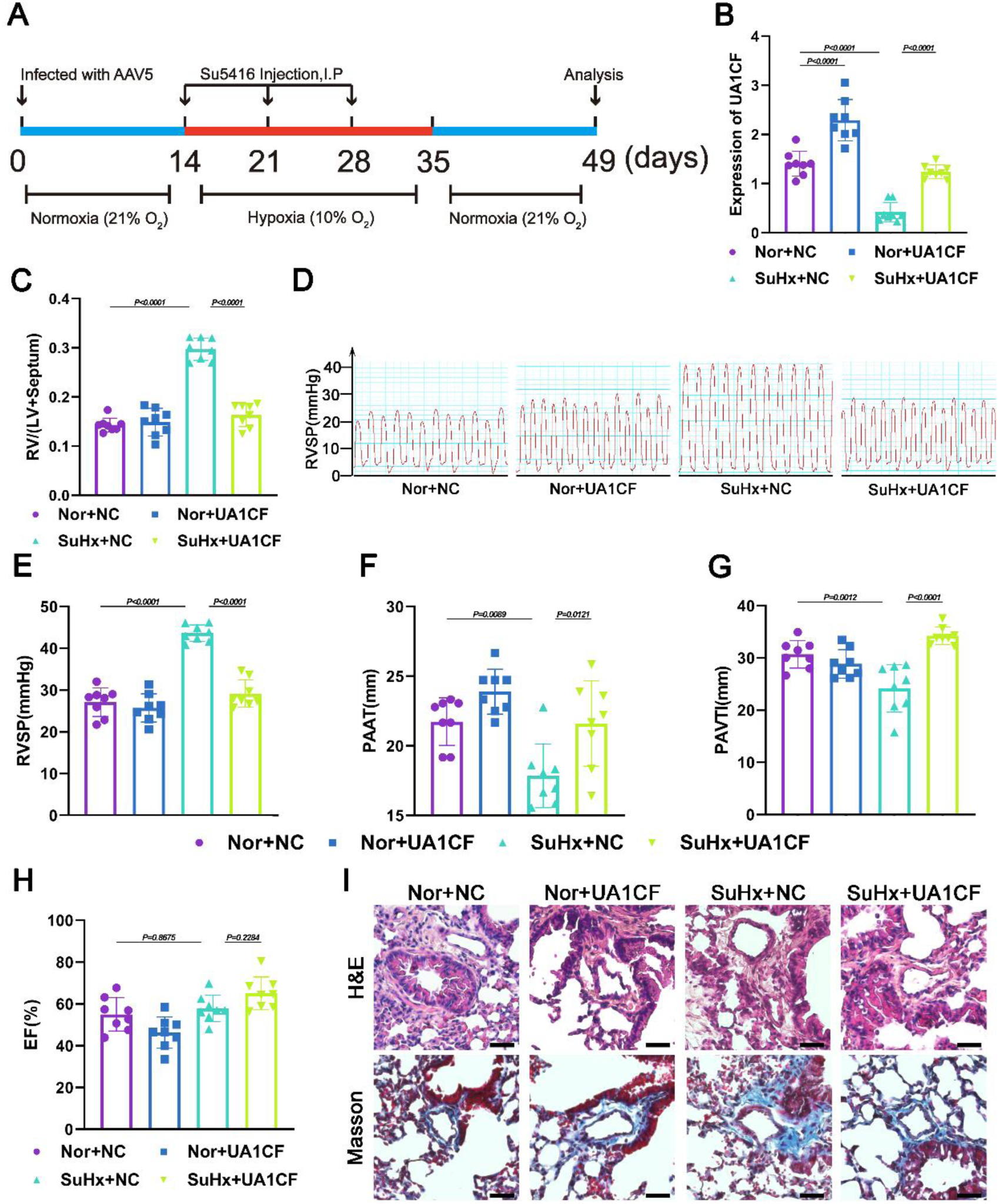
UNC5B-AS1 conserved fragment alleviates SuHx-induced PH in mice. **A** Schematic diagram of the SU5416 combined with hypoxia-induced PH (SuHx) mice model. **B** Expression of UNC5B-AS1 conserved fragment (UA1CF) in mice lungs (n=8). **C** Right ventricle/left ventricle + septum (RV/LV+ Septum) in mice (n=8). **D** Representative waveform of right ventricular systolic pressure. **E** Representative right ventricular systolic pressure (RVSP) in mice (n=8). Pulmonary artery acceleration time (PAAT, **F**), pulmonary artery velocity-time integral (PAVTI, **G**), ejection fraction (EF, **H**) in mice (n=8). **I** Representative images of Hematoxylin and Eosin staining (H&E) and Masson staining in mice lungs. Scale bars=50 μm. All data are presented as mean ± SEM. One-way ANOVA with Tukey post hoc test was used to compare multiple groups with equal variance, and Brown-Forsythe and Welch ANOVA with Tamhane T2 post hoc test was used to compare multiple groups with unequal variance. Nor, normoxia, SuHx, SU5416 combined with hypoxia, NC, Negative control and UA1CF, AAV5-UNC5B-AS1 conserved fragment overexpression.

### The UNC5B-AS1 conserved fragment alleviates pulmonary inflammation in PH mice

Next, we evaluated the impact of UA1CF on pulmonary lactate levels and the inflammatory response in mice. UA1CF overexpression significantly alleviated the decrease in mitochondrial complex IV activity (Fig.S7A) and the increases in LDH activity and lactate levels (Fig.S7B-C) in the lung tissues of SuHx mice. Immunofluorescence staining of lung tissues showed that UA1CF overexpression alleviated pulmonary arterial lactylation (Fig.S7D) and IL-6 expression (Fig.S7E) in SuHx mice. ELISA showed that UA1CF overexpression reduced the secretion of IL-1β, IL-6, and TNF-α in the lung tissues of SuHx mice (Fig.S7F). These results indicate that UA1CF can alleviate the pulmonary inflammatory response in SuHx mice.

## Discussion

Previous studies have shown that UNC5B antisense lncRNA 1 (UNC5B-AS1) is involved in and functionally relevant to a spectrum of biological events. This factor regulates signaling pathways by acting as an RNA decoy and microRNA sponge^33^, recruiting chromatin modifiers^34^ and inhibiting translation^35^. Recent studies have shown the involvement of UNC5B-AS1 in various cancers, and it can influence the growth and metastasis of human colon cancer cells by adsorbing miR-622 and ovarian cancer progression by regulating H3K27me on NDRG2 via EZH2^34^. However, the role of UNC5B-AS1 in the cardiovascular or respiratory systems, particularly in the pathogenesis of pulmonary arterial hypertension, remains unknown. The present study comprehensively screened super enhancer (SE)-associated lncRNAs in PH for the first time and showed that UNC5B-AS1 was driven by SEs in PASMCs. RNA-seq data showed that UNC5B-AS1 was widely downregulated in PASMCs under hypoxic conditions, while ChIP-seq and ChIP-qPCR assays identified UNC5B-AS1 as a super enhancer transcriptional target that was directly occupied by both H3K27ac and H3K4me1. This result was further verified by the finding that JQ1 (the BET bromodomain inhibitor) dramatically suppressed the expression of UNC5B-AS1. Based on our findings, the identified landscape of SEs may provide a better mechanistic understanding of the pathophysiological role of SE-lncRNAs in PH.

SEs affect gene expression programs through the binding of transcription factors and interactions with promoters^36^. To examine the precise factors that regulate UNC5B-AS1 expression in response to hypoxia, we analyzed the promoter of UNC5B-AS1 and found that there were multiple binding sites for transcription factors, such as FOXP3, YY1, and USF2. We observed that FOXP3 interacted with H3K27ac and H3K4me1 to maintain chromatin looping between the UNC5B-AS1 promoter and its super enhancers and contributed to its transcriptional activity in PASMCs. Although our data indicated that UNC5B-AS1 was regulated by hypoxia in a FOXP3-dependent manner, whether other transcription factors and epigenetic regulatory factors are involved in manipulating UNC5B-AS1 expression during hypoxic conditions in PASMCs requires further analysis.

Lactylation can occur in various cellular contexts, and is particularly under low oxygen or hypoxic conditions, catalyzed by lactate dehydrogenase enzymes^37^. Protein lactylation plays a crucial role in diverse biological processes, including energy metabolism^38^, cell signaling^39^, gene expression regulation^40^, and cell growth^41^. For instance, in tumor-infiltrating myeloid cells (TIMs), lactylation-driven METTL3-mediated RNA m6A modification enhances the immunosuppressive capacity of TIMs^39^. Furthermore, in myocardial infarction, histone lactylation regulates the dual activities of monocyte-derived macrophages by promoting the transcription of the repair genes Lrg1, Vegf-α, and IL-10, thereby enabling anti-inflammatory and proangiogenic responses^41^. Our research is the first report on the key role of lactylation in the occurrence and development of PH. The SE-lncRNA UNC5B-AS1 plays a crucial role in maintaining the contractile phenotype of PASMCs. Under hypoxic conditions, the loss of UNC5B-AS1 decreases the binding of LRPPRC to mitochondrial complex IV, impairing its activity and leading to aberrant activation of glycolysis, which ultimately causes intracellular lactate accumulation in PASMCs. Excessive lactate in PASMCs leads to lactylation of the upstream promoter regions of the IL-1β, IL-6, and TNF-α genes, thereby promoting the transition of PASMCs toward an inflammatory phenotype. This results in the release of inflammatory factors that participate in the vascular perivascular inflammatory microenvironment. By overexpressing UNC5B-AS1 in cultured PASMCs, we further demonstrated that the hypoxia-induced inflammatory phenotype in PASMCs and the release of inflammatory factors occur in a UNC5B-AS1-dependent manner. These results indicate that UNC5B-AS1 plays an important role in the development of pulmonary hypertension. Studying the upstream lactylation sites on key genes and investigating the functions of their respective writers, readers, and erasers in PH is important.

The interaction between lncRNAs and proteins has been shown to play an important role in various biological processes. It has been reported that LRPPRC forms an RNA-dependent protein complex that is necessary for maintaining a pool of nontranslated mRNAs in mammalian mitochondria^42^. Studies have shown that the fibroblasts and skeletal muscle homogenates of mitochondrial disease patients showed decreased protein levels of LRPPRC and impaired complex IV enzyme activity, which were associated with abnormal complex assembly and reduced steady-state levels of numerous oxidative phosphorylation subunits^43^. The lncRNA SNHG17 has been shown to stabilize c-Myc protein and promote G1/S transition and cell proliferation by interacting with LRPPRC^44^. It can be hypothesized that LRPPRC is crucial for maintaining mitochondrial oxidative respiratory chain activity and is capable of binding to lncRNAs to exert biological effect. Our present study showed that UNC5B-AS1 could act as an LRPPRC molecular scaffold, thereby interfering with mitochondrial complex IV to modulate oxidative phosphorylation. These findings suggest that UNC5B-AS1 regulates glycolysis by modulating LRPPRC and mitochondrial complex IV. Ultimately, through lactylation, UNC5B-AS1 affects the perivascular inflammatory microenvironment and the inflammatory phenotypic transition of PASMCs, indicating that UNC5B-AS1 overexpression may provide therapeutic options for PH.

The highly conserved region of UNC5B-AS1 that binds to LRPPRC functional fragment was identified using bioinformatics tools in the mouse genome. We found that the expression of the UNC5B-AS1 conserved fragment named UA1CF was absent in the lung tissues of PH mice induced by SU5416 combined with hypoxia (SuHx). Additionally, in mouse PASMCs, UA1CF could bind to LRPPRC. Adenovirus infectionwas used to overexpress UA1CF in the lungs of mice. We discovered that UA1CF could alleviate the symptoms of PH induced by SuHx in mice. Similarly, UA1CF expression in mouse lung tissues influenced the activity of mitochondrial complex IV, as well as the lactate levels and the extent of panlactylation modification in the lungs. Finally, we evaluated the levels of the inflammatory factors IL-1β, IL-6, and TNF-α, in mouse lung tissues and found that UA1CF significantly alleviated pulmonary inflammation in PH mice. Based on these results, we confirmed that UA1CF could alleviate the symptoms of PH and the levels of pulmonary inflammation in SuHx mice and provided substantial evidence supporting the beneficial role of UNC5B-AS1 in PH. Furthermore, it would be interesting to further investigate whether UA1CF was part of another novel lncRNA in the mouse genome.

In conclusion, the present study demonstrated that the SE-driven lncRNA UNC5B-AS1 plays a vital role in alleviating the inflammatory phenotype of PASMCs, thereby regulating the pathological remodeling of vessels in PH. Mechanistically, UNC5B-AS1 functioned as a novel checkpoint molecule in oxidative phosphorylation by acting as an LRPPRC molecular scaffold, thereby modulating the activity of mitochondrial complex IV. Downregulation of UNC5B-AS1 reduced the binding of LRPPRC to mitochondrial complex IV, leading to increased lactic acid levels and promoting lactylation of the promoter regions of IL-1β, IL-6, and TNF-α, thereby contributing to a perivascular inflammatory microenvironment that stimulates the proliferation of surrounding PASMCs. This finding implicates SE-lncRNAs in PH and suggests that the modulation of UNC5B-AS1 activity and its downstream targets, such as LRPPRC, is a novel therapeutic approach for treating this hypoxia-associated disease.

## Acknowledgments

X.Z. designed the research, performed the experiments, analyzed the results, and wrote the manuscript. X.Z., X.W., X.G., L.Z., Z.W., X.P., J.M., L.O., Z.M., and Y.L. performed the experiments and analyzed the results. X.W., and C.M. performed and analyzed the RNA sequencing and ChIP sequencing, supervised the research, and wrote the manuscript. X.Z., and Y.C. supervised the study. F.L. and X.Z. performed the animal experiments and interpreted the results. C.M. designed and supervised the research and approved the final version of the manuscript.

## Supplemental Material

Supplemental Methods

Tables S1-S4

Figure S1-S7

## Non-standard Abbreviations and Acronyms

UNC5B-AS1: UNC5B antisense RNA 1
PASMCs: Pulmonary Artery Smooth Muscle Cells
PH: Pulmonary Hypertension
lncRNA: Long Noncoding RNA
SE: Super Enhancer
ChIP: Chromatin Immunoprecipitation
LRPPRC: Leucine-Rich PPR Motif-Containing Protein
FOXP3: Forkhead Box Protein P3
IL: Interleukin
TNF-α: Tumor Necrosis Factor Alpha
OPN: Osteopontin
KLF4: Krüppel-Like Factor 4
CNN1: Calponin-1
SM22α: Smooth Muscle Protein 22-alpha
HKII: Hexokinase II
PKM2: Pyruvate Kinase M2
ECAR: Extracellular Acidification Rate
CM: Conditioned Medium
SuHx: Sugen5416/Hypoxia
AAV5: 5 Adenovirus-Associated Virus
H&E: Hematoxylin And Eosin Staining
UA1CF: UNC5B-AS1 Conserved Fragment
RV/LV: Right Ventricle/Left Ventricle
RVSP: Right Ventricular Systolic Pressure
PAAT: Pulmonary Artery Acceleration Time
PAVTI: Pulmonary Artery Velocity-Time Integral
EF: Ejection Fraction

